# Reinforced CRISPR interference enables reliable multiplex gene repression in phylogenetically distant bacteria

**DOI:** 10.1101/2025.04.14.648646

**Authors:** Joshua R. Elmore, Ritu Shrestha, Andrew Wilson, Elise Van Fossen, Andrew Frank, Ryan M. Francis, Henri Baldino, Molly Stephenson, Bhavya Gupta, Jeane Rivera, Robert G. Egbert

## Abstract

Genetic screens are essential for uncovering novel molecular mechanisms and identifying the functions of hypothetical proteins. CRISPR interference (CRISPRi) is a powerful, programmable, and sequence-specific gene repression technology that can be used for high-throughput screening and targeted gene repression. Despite its ease of use, the initial development of CRISPRi systems is labor-intensive in many non-model organisms. Our goal is to simplify this by establishing a host-agnostic CRISPRi platform that utilizes the serine recombinase-assisted genome engineering (SAGE) system. This system integrates CRISPRi machinery directly into the bacterial chromosome, overcoming the limitations of plasmid-based systems and enabling wide sharing across diverse bacteria. We demonstrate the design and optimization of multiplexed CRISPRi to repress multiple genes simultaneously in phylogenetically distant bacteria. We use a *Francisella novicida*-derived Cas12a system that processes multiple distinct CRISPR RNAs, each targeting a unique gene sequence, from a single transcript. This allows easy multi-gene repression. By reinforcing gene repression with multiple guides targeting a single gene, we achieve robust genetic perturbations without the need to pre-screen the efficacy of guide RNAs. Using this toolkit, we perturb multiple combinations of growth and visual phenotypes in Pseudomonas fluorescens and demonstrate simultaneous repression of multiple fluorescent proteins to near background levels in bacteria from various other genera. While the tools are directly portable to all SAGE-compatible microbes, we illustrate the utility of SAGE by optimizing CRISPRi performance in *Rhodococcus jostii* through a combinatorial screen of Cas protein and CRISPR array expression variants. The efficient integration of CRISPRi machinery via the SAGE system paves the way for versatile genetic screening, enabling profound insights into gene functions both in laboratory conditions and relevant naturalistic scenarios.

## Introduction

Genetic screens are a critical tool for discovering novel molecular mechanisms and identifying the functions of hypothetical proteins – which frequently comprise ∼40-60% of sequenced bacterial genomes (*1–4*). CRISPR interference (CRISPRi) is a programmable, sequence-specific gene repression technology that can be used for both high-throughput genetic screens and targeted gene repression (*5–9*). CRISPRi works by using a guide RNA (crRNA or sgRNA) to direct a catalytically inactive Cas protein to bind a specific DNA sequence (**Fig. 1** – diagram of CRISPRi function). Once bound, the protein-RNA complex (effector) blocks RNA polymerase from completing transcript elongation, resulting in sequence-specific gene repression. While the ease of employing CRISPRi systems has led to its wide-spread adoption for evaluating gene functions, their initial development takes substantial effort and the systems typically only function in a narrow range of related organisms. Here we created highly effective CRISPRi systems in phylogenetically diverse bacteria by developing a host-agnostic CRISPRi development platform.

**Fig 1.**
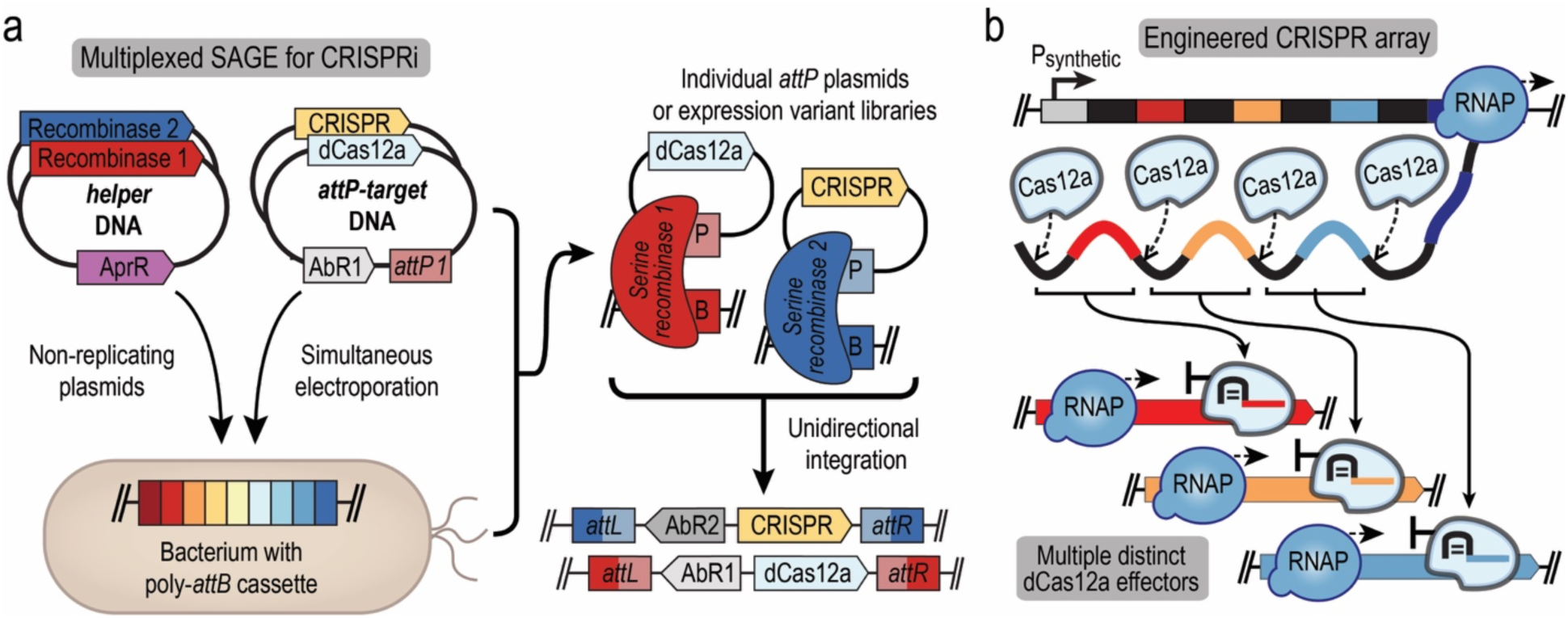
Simultaneous SAGE integration of CRISPRi machinery for multiplex gene repression. (a) Simultaneous SAGE integration of two DNA constructs for expression of a DNase-deactivated Cas12a variant (dCas12a) and expression of an engineered CRISPR array. Constructs are transferred into host organisms that have been engineered with a poly-*attB* cassette that serves as a ‘landing pad’ for site-specific chromosomal integration of DNA. Each SAGE recombinase is transiently expressed from a nonreplicating helper plasmid, catalyzing recombination between its cognate *attP* and *attB* sequences, located on the co-transferred target plasmid and the host chromosome, respectively. (b) Primary CRISPR array transcript processing by dCas12a produces distinct effector complexes that repress transcription of target genetic loci by disrupting RNA polymerase (RNAP).

Advances in functional genomics have enabled the determination of certain classes of proteins (e.g., biosynthetic enzymes), but most genes in environmental bacteria do not have an identified phenotype under laboratory conditions (*3, 4*). Nearly all functional genomics experiments have been restricted to purely laboratory conditions and do not interrogate the importance of genes under relevant and diverse naturalistic conditions. Experiments must be performed in conditions that better reflect natural environments to identify the functions of these genes. However, common methods used to deliver and control CRISPRi systems in bacteria have substantial caveats that limit their utility outside of controlled laboratory environments. Most bacterial CRISPRi systems are maintained in the host using autonomously replicating plasmids (*7, 10–14*) and are often designed to function in a narrow range of bacterial hosts. Most replicating plasmids, including the most commonly broad-host range variants (e.g., pBC1 (*15*), pBBR1 (*16, 17*), RK2 (*18, 19*), RSF1010 (*20*)) are generally unstable, often reduce the fitness of the host, and require constant selection – typically with antibiotic resistance (*21–28*). A commonly touted advantage of CRISPRi over other genetic screening methods (e.g. genome-wide transposon insertion mutant sequencing) is that repression can be tuned by titrating guide RNA and Cas protein expression (*5, 8*). However, the addition of exogenous compounds to maintain plasmids and control expression is both impractical in most naturalistic environments and alters native conditions. Furthermore, replicating plasmid copy number is highly variable both within a species (*29–31*) and between species (*32, 33*), thus adding cell-cell variability to CRISPRi phenotypes, including the selection of CRISPRi-defective mutants. Varying gene dosage affects the extent of gene repression (*6*) and Cas-protein associated toxicity (*17, 34, 35*) thus harboring CRISPRi machinery on variable copy plasmids complicates the interpretation of genetic screen results.

The majority of bacterial CRISPRi systems use deactivated Cas9 (dCas9) with a single guide RNA (sgRNA) (*5*) and Cas9-based systems are almost exclusively designed for single gene repression. In dCas9- based systems, each sgRNA must be transcribed independently with its own transcriptional promoter and terminator sequences. Recent work has begun to address the technical complexity of constructing genetically stable multi-sgRNA expression constructs (*36*) with distinct promoters and terminators. A requirement to identify a family of sequence-distinct and effective promoter/terminator pairings renders effective sgRNA design for many non-model hosts inaccessible. Shared promoters often function differently between organisms and terminators are poorly characterized in most bacteria.

Though used less frequently than Cas9 for CRISPR interference, DNase-deactivated variants of Cas12a (dCas12a) have features that simplify multiplexed gene repression (*37–39*). In addition to cleaving DNA, the Cas12a enzyme processes CRISPR array transcripts that contain multiple distinct guide sequences into individual CRISPR RNAs (crRNAs) (Figure 1) (*39*). The ability to place multiple guides into a single and highly compact DNA construct greatly simplifies design, construction, and optimization of dCas12a-based CRISPRi systems. Additionally, this transcript processing makes robust transcriptional termination less critical than with transcription of the sgRNAs used in dCas9-based systems.

A host-agnostic CRISPRi platform must simultaneously repress multiple genetic loci, reliably operate despite ineffective individual guide RNAs, stably function without antibiotic selection, and easily extend to phylogenetically diverse bacteria. Expression of both the Cas protein and guide RNAs must be fine-tuned to ensure robust gene repression without reducing cell fitness due to excessive Cas protein expression (*17, 34, 35*). Most challenges associated with applying CRISPRi in naturalistic environments across a broad range of host organisms can be solved by constitutively expressing host-optimized CRISPRi machinery from the host chromosome. Multiple bacteriophage recombinase-based technologies, such as SAGE and CRAGE, can be used efficiently incorporate genetic payloads into bacterial chromosomes (*28, 40*).

Here we use SAGE (<u>S</u>erine recombinase-<u>A</u>ssisted <u>G</u>enome <u>E</u>ngineering) (*28*) to efficiently incorporate CRISPRi machinery into the chromosomes of phylogenetically distant bacteria and fine-tune its expression. SAGE draws from multiple site-specific recombinases encoded on non-replicating plasmids to unidirectionally integrate up to 10 DNA constructs into the bacterial chromosome at high efficiency. (**Fig. 1a**). These features enable SAGE to be used in virtually any bacterium. Our CRISPRi platform utilizes the dCas12a enzyme from *Franciselle novicida* (*39*) and leverages the simplicity of dCas12a RNA processing to reinforce repression by encoding multiple guide RNAs for each targeted gene. We use the high efficiency of SAGE (*28*) to simultaneously incorporate both the dCas12a encoding gene and a collection of CRISPR arrays into the host chromosome (**Fig. 1a**). This enables rapid CRISPRi optimization across a phylogenetically diverse range of bacteria, which we have demonstrated for strains of *Pseudomonas*, *Rhodococcus*, and *Corynebacterium* used for plant growth promotion, bioremediation, and industrial biomanufacturing applications.

## Results

### Rapid optimization of Cas12a expression from the chromosome

We initially demonstrated our CRISPRi platform on the model plant growth-promotion rhizobacterium *Pseudomonas fluorescens* SBW25. Since bacterial fluorescence is a simple and effective measurement for both bulk populations and single cells, we evaluated repression of sfGFP to optimize expression of the CRISPRi machinery. We constructed strain JE4694, a genetic variant of SBW25 that harbors both a poly-*attB* cassette that is used for SAGE integration (*28*) and a highly expressed sfGFP-*cat* (chloramphenicol acetyltransferase) translational fusion gene. For simplicity we will refer to the sfGFP-*cat* fusion protein as sfGFP. All SBW25 and other host strains containing CRISPRi machinery were generated by simultaneously incorporating an *attP*-target plasmid containing a CRISPR array into the Bxb1 *attB* site and a second *attP*-target plasmid containing a dCas12a expression cassette into the TG1 *attB* site (**Fig. 1a**). The CRISPR arrays are modeled on native *Fn* Cas12a-associated CRISPR arrays (*39*), replacing the native upstream region with synthetic promoters (**Fig. 1b**).

We optimized transcription and translation of dCas12a to minimize CRISPRi toxicity and to maximize gene repression (**Fig. 2a**). Multiple studies have documented reduced growth rates associated with the excessive expression of deactivated Cas proteins in organisms (*17, 34, 35*). We generated a pooled combinatorial library of dCas12a expression variants for integration into SBW25 that contain a collection of ribosomal binding sites (RBSs) and transcriptional promoters. RBS sequence variation was predicted to modulate translation over a 25-fold range (*41*) and promoter sequence variation spans a 45-fold range of expression in *Pseudomonas putida* KT2440 (*42*) (**Supplementary Table T3**). The dCas12a expression library and a CRISPR array that expresses a single sfGFP-targeting guide called CmR were simultaneously integrated into the JE4694 chromosome, and the pooled strain library was evaluated for sfGFP repression.

**Fig 2.**
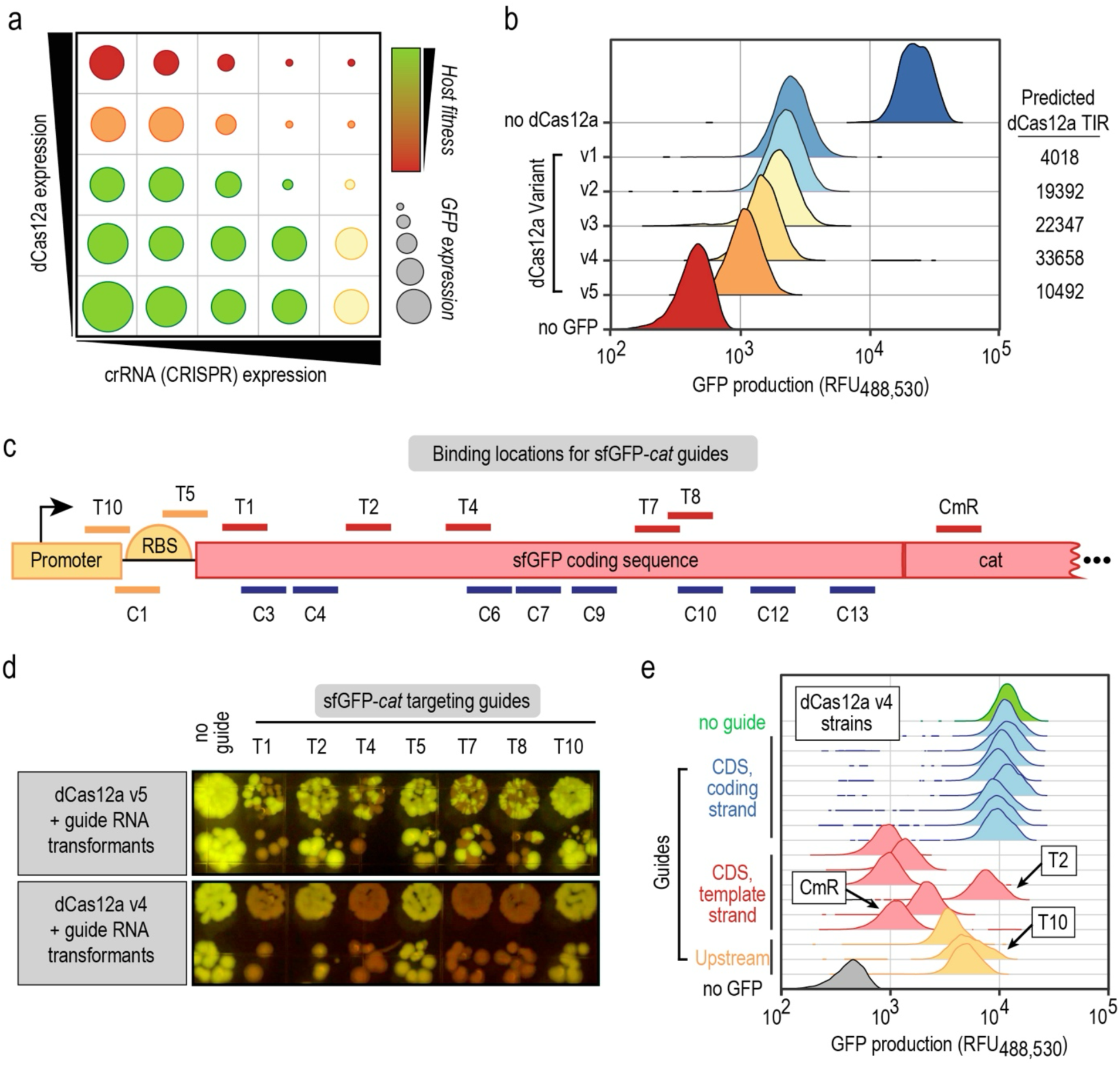
Developing a CRISPR interference system in *P. fluorescens* SBW25 and identifying guide targeting rules. (a) Conceptual graphic demonstrating the theoretical relationship between CRISPR & Cas protein expression with the efficiency and toxicity of CRISPR interference. Increasing toxicity is represented by low fitness (red) and high fitness (green) and the efficacy GFP repression is indicated by increasingly small circles. Ideal CRISPRi is represented by green color and small circle size. (b) Representative flow cytometry measurements of *P. fluorescens* SBW25 variants that either do not express sfGFP (no GFP), express GFP but lack CRISPRi machinery (no dCas12a), or express the sfGFP-CmR (GFP), a CmR guide and varying levels of dCas12a (dCas12a variants). (c) Diagram of sfGFP-CmR and the sites targeted by each guide. (d) Images of SBW25 transformants that express the indicated dCas12a variant (left) and contain a CRISPR array with the composition indicated above. (e) Flow cytometry measurements of SBW25 strains that do not express sfGFP (no GFP), express GFP but lack CRISPRi machinery (no guide), or express the sfGFP, harbor dCas12a v4, and express the indicated guide variant. (b,e) The *x* axis indicates the relative fluorescence (excitation at 488 nm and emission at 530 nm) of each cell. The *y* axis represents the abundance of cells in the population with a given relative fluorescence unit (RFU).

We identified multiple CRISPRi expression variants from the pooled library that effectively repress sfGFP expression. Transformant colonies with robust sfGFP repression were substantially darker than other library members when visualized on a blue light box. We identified which expression variants robustly repressed sfGFP by sequencing the promoters and RBSs in 10 of the darkest transformants. Five RBSs were represented in the 10 strains and with only two exceptions, each strain used the strongest of the four promoters to drive expression of dCas12a (**Supplementary Table T3**). Of note, none of the strains used the predicted weakest promoter, PJE121211, to drive dCas12a expression. Expression variants that best repressed sfGFP were identified by constructing dCas12a expression variants with one the five identified RBS sequences and the strongest of the four dCas12a promoters (Ptac) and evaluating the resulting strains for their ability to repress sfGFP expression (**Fig. 2b**). Apart from one expression variant, the repression of sfGFP scaled with the predicted dCas12a translation initiation rate (TIR). Interestingly, the highest sfGFP repression (30.5-fold) was associated with a variant with the second lowest predicted TIR (variant v5), likely reflecting an unmodeled factor that increases translational efficiency.

### Repression is dependent upon the strand and position in the gene bound by the dCas12a effector

The efficacy of CRISPRi can be dependent upon which strand of DNA and what position within the gene (e.g., promoter, coding region) is bound by the CRISPR/Cas effector (*43*). Our initial CmR guide targets the template strand within the middle of the sfGFP-*cat* gene and supported robust repression of sfGFP. So, we next evaluated whether strand (template or coding) or position of target has an impact upon repression of sfGFP. A collection of 17 dCas12a effectors whose guides target unique sequences within the sfGFP*-cat* gene (**Fig. 2c**) were evaluated for their ability to block sfGFP expression. Of these, 3 guides target the region upstream of the start codon (**Fig. 2c, orange**) and 15 target the coding sequence (CDS) on either the template (**Fig. 2c, magenta**) or coding (**Fig. 2c, blue**) strand. We initially assessed performance of strains using dCas12a expression variant v5 and observed substantial variability in colony morphology and fluorescence between transformants with the same DNA constructs (**Fig. 2d, Supplementary Fig. S1**). We posited that toxicity arising from excessive dCas12a expression led to genetic instability. So, we used the more reliable v4 expression variant (**Fig. 2b-c**) to evaluate the impact of target sites on gene repression.

We observed two general trends in sfGFP repression (**Fig. 2e**). First, targeting the template strand of the CDS elicits a stronger sfGFP repression phenotype than targeting upstream non-coding sequences and position within the CDS had no apparent influence on repression. Second, targeting the coding strain of CDS does not elicit a substantial sfGFP repression phenotype. While five of the six template strand-targeting guides elicit between 6.2 to 24.6-fold sfGFP repression, one guide (T2) failed to elicit a substantial repression phenotype.

### Reinforcing CRISPRi with multiple guides per gene improves repression multiplicatively

The inconsistency of gene repression on a guide-to-guide basis and the potential need for very strong repression to elicit a phenotype necessitates improved and more reliable CRISPR interference. As highlighted in the genetic instability of the most highly expressed Cas variant (**Fig. 2d**), increasing expression of Cas or CRISPR components is not a viable solution. Gene repression is a consequence of dCas12a effectors physically blocking transcription by RNA polymerase (RNAP) (**Fig. 3a-b**) and thus transcription only occurs when the Cas effector complex is disassociated from the DNA or when it is physically displaced by RNA polymerases. So, rather than increasing the number of effectors targeting the same position on the gene by increased expression, we attempted to target multiple effector complexes across the gene, blocking transcription by RNAPs that ‘escape’ the initial dCas12a effector (**Fig. 3a-b**) thereby enhancing gene repression without changing CRISPRi machinery expression levels.

**Fig. 3.**
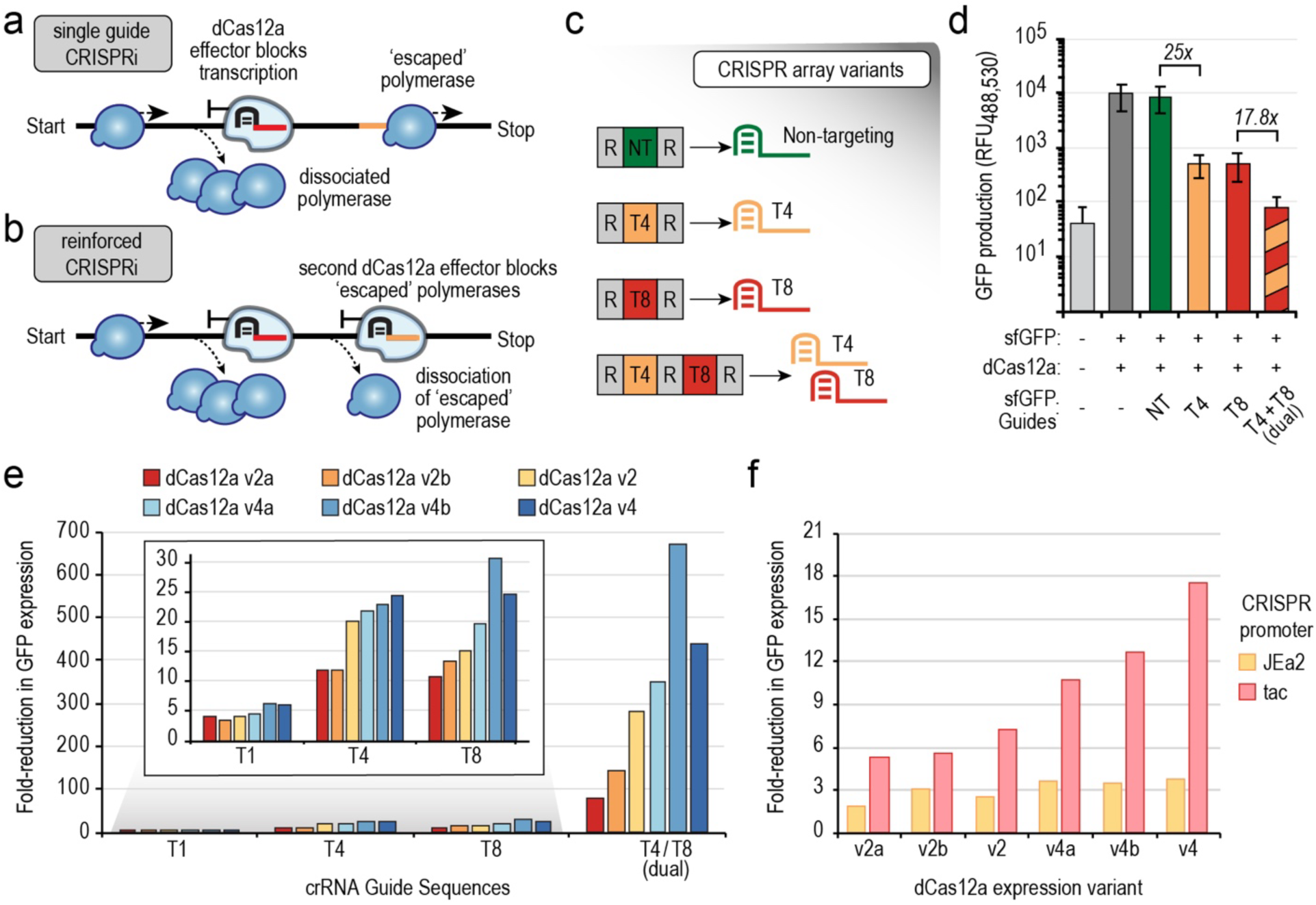
Enhancing the robustness of CRISPR interference. (a-b) Graphical representation of how dCas12a effector complexes prevent gene expression by physically blocking mRNA extension by RNA polymerase (blue). Reinforced CRISPRi reduces gene expression by blocking RNA polymerases that bypass an initial effector complex (a) with a second effector complex that binds a downstream site in the gene (b). (c) Schematics of CRISPR arrays used to assess how targeting GFP with one or more guides that bind either one or two sites in the gene affect repression in repression of GFP expression in panel d. (d) Graph of GFP production in P. fluorescens SBW25 strains with the presence of sfGFP, dCas12a, and composition of CRISPR array (if present) is listed below. Y-axis represents the mean fluorescence as determined by flow cytometry. Error bars represent two-sided standard deviations of the median values from four biological replicates. (e-f) Graphs indicate extent to which GFP expression is repressed in strains with (e) dCas12a expression or (f) CRISPR array expression variants. Values represent the ratio of median fluorescence of four biological replicates from in a strain without a CRISPRi system versus a strain containing the indicated CRISPRi system and guide sequence(s).

Similar to prior studies in *E. coli* (*44*), we found that targeting a gene with multiple guides greatly enhances gene repression. We evaluated whether targeting dCas12a effectors to two distinct positions in a CDS would enhance gene repression by measuring sfGFP expression when using CRISPR arrays that include either a one or two sfGFP-targeting guides (**Fig. 3c-d**). We observed substantially 25-fold reduction in sfGFP expression when using either guide alone, but when both are guides are simultaneously expressed from a single CRISPR array we observed a further 17.8-fold reduction (437-fold overall reduction). This confirms that targeting multiple positions on a gene, which we call reinforced CRISPRi greatly increases gene repression without altering CRISPRi machinery expression.

### Optimizing CRISPR/dCas12a expression to maximize repression and minimize the burden of CRISPRi

Next, we systematically assessed to what extent we could further reduce dCas12a and CRISPR array expression without affecting gene repression by evaluating performance of a small combinatorial collection of dCas12a and CRISPR array expression variants (**Fig. 3e-f**). Each dCas12a variant was also evaluated with four different sets of guides to determine how consistence repression is when using distinct guides. Specifically, we evaluated six dCas12a variants that combine three of the four promoter variants from our original library that span a predicted 7.2-fold range of expression (PJE111111, PJE121111, and PJE111211) (*42*) with the RBSs from v2 and v4 dCas12a expression variants (**Fig. 2b**).

The expression of sfGFP largely decreases proportionally with predicted increases in dCas12a expression. However, we observed one exception. The v4b expression variant, which combines the PJE11121 promoter with the v4 RBS (**Fig. 3e**), performs similarly to the original v4 variant despite using a significantly weaker promoter (7-fold in Elmore, *et al*) (*42*). This suggests the amount of dCas12a protein produced by the v4 variant is well in excess of what required to maximize CRISPRi activity and thus can be substantially reduced to lower use of cellular resources.

The influence of CRISPR array expression was evaluated using two distinct promoters, P*tac* and PJEa2, the former of which is used for most experiments in this study (**Fig. 3f**). The activities of these promoters are consistent across conditions in *P. fluorescens* SBW25 and span a 11.6-fold range of expression (*28*). We found that expression of a CRISPR array containing the CmR guide is saturated with the *tac* promoter, as sfGFP expression is dependent upon dCas12a expression levels. However, the opposite appears to be true with the PJEa2 promoter, where sfGFP expression is independent of dCas12a expression, suggesting that availability of the crRNA is limiting. These findings suggest that expression of both dCas12a and the CRISPR array can be reduced while maintaining the full efficacy of CRISPRi, and that the expression of these components is interdependent.

### Simultaneous repression of multiple genes using reinforced CRISPRi

We evaluated repression of three fluorescent protein (FP) encoding genes to assess whether reinforced CRISPRi can support multiplex gene repression when guide sequences are randomly chosen. For this, we chose the fluorescent proteins sfGFP (green), mKate2 (far red), and mOrange2 (orange) because they are bright and have minimal spectral overlap. For these, and all further experiments in *P. fluorescens* we utilize Ptac for CRISPR array expression and the Cas12a v4b expression variant. Initially, we evaluated simultaneous repression of sfGFP and mKate2. For this, we used CRISPR arrays that each have two guides for sfGFP and either a single or combination of guides that target mKate2 (**Fig. 4a**) in strain JE4807, which expresses both proteins at high levels (**Fig. 4b**). In parallel, we performed analogous experiments in strain JE4815 which expresses sfGFP and mOrange2 at high levels. For these experiments we used CRISPR arrays containing guides targeting all three fluorescent proteins (**Fig. 4a**). Additionally, we used control CRISPR arrays that either only contain sfGFP-targeting guides or contain no gene-targeting guides.

**Fig. 4.**
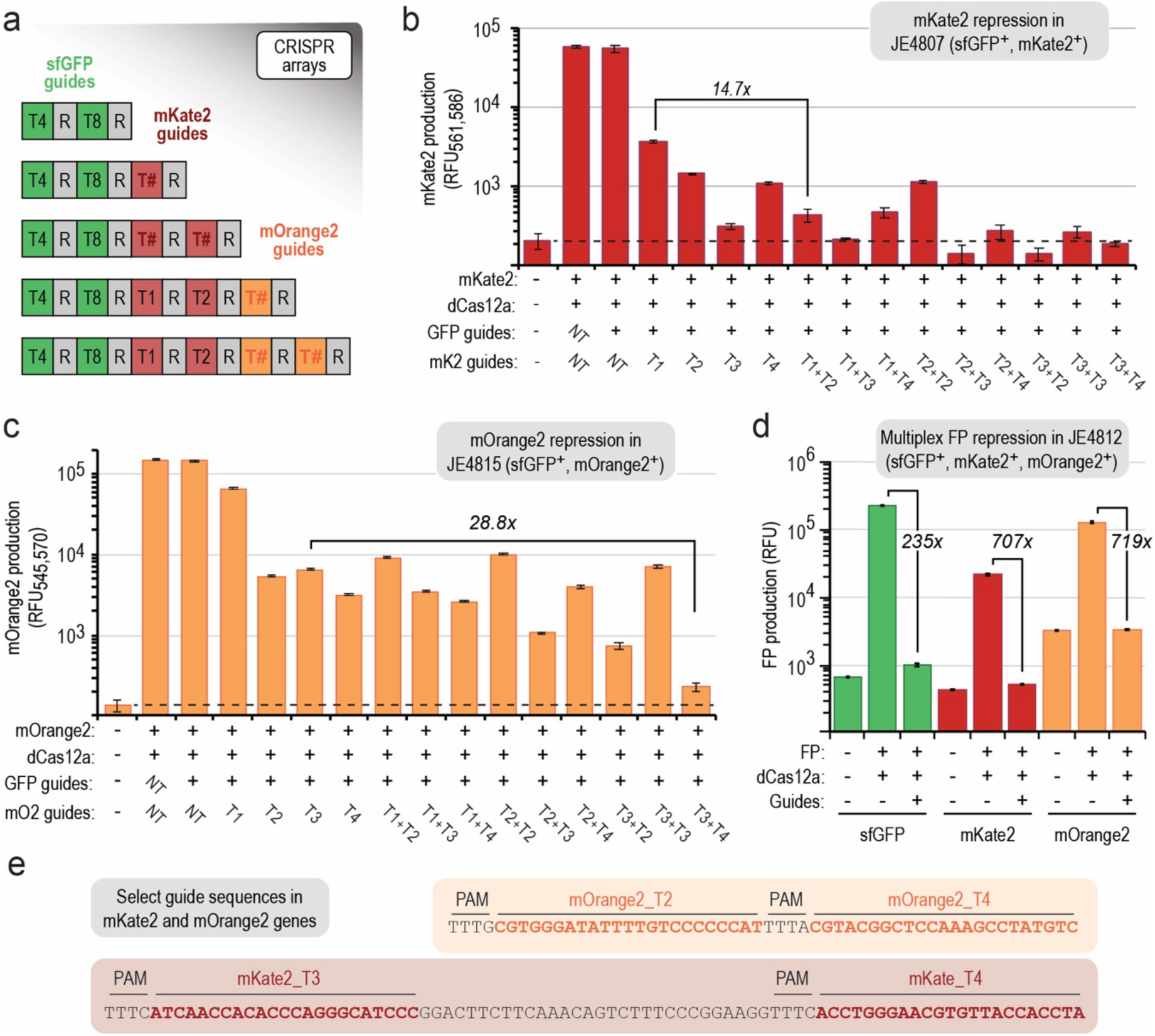
Reinforced CRISPRi enables robust multiplex gene repression. (a) Schematic of CRISPR arrays used for multiplexed repression of green, red, and orange fluorescent proteins in panels b-d. T# values represent T1, T2, T3, or T4. (b-c) Graph displaying production of (b) mKate2, (c) mOrange2, or (d) all three fluorescent proteins in *P. fluorescens* SBW25 strains that express either (b) sfGFP and mKate2 or (c) sfGFP and mOrange2 or all three proteins (d). (b-d) Presence of fluorescent protein, dCas12a, and composition of CRISPR arrays are listed below. Y-axis values represent either (b) the mean fluorescence as determined using flow cytometry or (c-d) plate reader measurements. Error bars represent the two-sided standard deviation in 4 biological replicates. (e) Sequences from the mOrange2 gene (top) or mKate2 gene (bottom). Sequences targeted by guides are bold and colored. Protospacer adjacent motifs (PAM) sequences are indicated.

Overall, multiplex repression of two fluorescent proteins was robust. As expected, strong repression of sfGFP was observed in strains with arrays containing the two sfGFP-targeting guides (**Supplementary Fig. S2**). With only a single exception (mOrange_T1), all arrays containing mKate2- or mOrange2- targeting guides enable robust repression of their respective FP-encoding genes (**Fig. 4b-c**). In most cases, reinforced CRISPRi greatly improved repression over standard CRISPRi, and doubling the copy number of the same guide had minimal impact on repression. Additionally, the three genes are simultaneously repressed by 235-fold to 719-fold when CRISPRi machinery targeting the three genes is expressed in JE4812 – a strain that expresses all three fluorescent proteins (**Fig. 4d**).

Two exceptions where the dual guide approach did not greatly improve gene repression highlight both the advantages and design constraints of reinforced CRISPRi. First, one guide failed to elicit substantial repression of mOrange (T1), so it did not contribute to improved repression when combined with other guides (**Fig. 4c**). While conventional CRISPRi would not elicit a phenotype using the T1 guide, reinforced CRISPRi enables repression of mOrange with the T1 by pairing the non-functional guide with a functional guide (T2-T4).

Using multiplicative increase in gene repression as a proxy for simultaneous effector binding on two distinct sites in a gene we interrogated how close effectors can be located before they become antagonistic. When target sequences were 34 bp apart in the mKate2 gene (**Fig. 4e**), the repression of mKate2 was substantially higher than observed with single guides (**Fig. 4b**). However, when target sequences were 4 bp apart in the mOrange2 gene (**Fig. 4e**), the repression of mOrange2 was similar to that or individual guides – suggesting that both effectors were unable to bind simultaneously. Interactions with sequence immediately upstream of the target, known as the protospacer adjacent motif (PAM), is critical for effector complex binding and this data suggests that placement of two targets immediately adjacent to each other prevents interactions with the PAM. While we did not identify the minimal distance required for reinforced CRISPRi guide placement, this experiment demonstrates that a 34 bp gap is sufficient for simultaneous binding of two effector complexes.

### Multiplex repression of diverse phenotypes using dCas12a-based CRISPR interference

As a practical demonstration, we next used reinforced CRISPRi to simultaneously repress multiple native and synthetic genes whose disruption will demonstrate visible phenotypes. These phenotypes include inability to grow when provided specific carbon and nitrogen sources, reduction of fluorescence, and mucoidy (see **Supplementary Fig. S3** for example). To prevent consumption of acetate and octanoic acid as carbon sources, we targeted repression of *aceA* (PFLU3817) or *glcB* (PFLU5623), which are predicted to encode isocitrate lyase and malate synthetase G, respectively. Blocking metabolic flux through these two enzymes prevents growth using carbon sources such as acetate and octanoate that require the glyoxylate shunt to biosynthesize C4 and larger central metabolites. To prevent of assimilation of specific nitrogen sources, we targeted *narB* (PFLU3426), *ureC* (PFLU0578), or *ntrC* (PFLU0343) which are predicted to encode nitrate reductase, the *α*-subunit of urease, and a transcriptional activator of nitrogen-limitation induced genes (e.g., *narB* and *ureC*). Disruption of *ureC* and *narB* expression should abolish or slow the assimilation of nitrate and urea as nitrogen sources. To generate a mucoid phenotype, we targeted *mucA* (PFLU1468), a which is predicted to encode a transcription factor that regulates exopolysaccharide (EPS) production.

We evaluated visual phenotypes displayed by strains containing dCas12a v4b and CRISPR arrays that either do not target genes in *P. fluorescens* genome or target combinations of the genes listed above (**Fig. 5a-c**). For this, we used a solid MOPS-based mineral medium (MME) (*45*) containing either sodium citrate, glucose, sodium acetate, or sodium octanoate as the carbon source and ammonium sulfate, urea, or sodium nitrate as the nitrogen source. All strains grew robustly when supplied the control carbon and nutrient sources glucose and ammonium. Aside the strain that harbors the non-targeting control array, each strain harbored guide sequences targeting *mucA*. Accordingly, we observed a mucoid phenotype for each of these strains when cultivated on MME containing glucose and ammonium sulfate (**Fig. 5b**). The mucoid phenotype complicates analysis of other growth phenotypes by both increasing OD per cell and changing colony morphology, so we used citrate, a carbon source where the mucoid phenotype was reduced (**Fig. 5c**), as the base carbon source for assays evaluating the other phenotypes. As expected, green fluorescence was substantially reduced in strains expressing crRNA containing guides that target sgGFP **(Fig. 5c – see bottom of panel).**

**Fig. 5.**
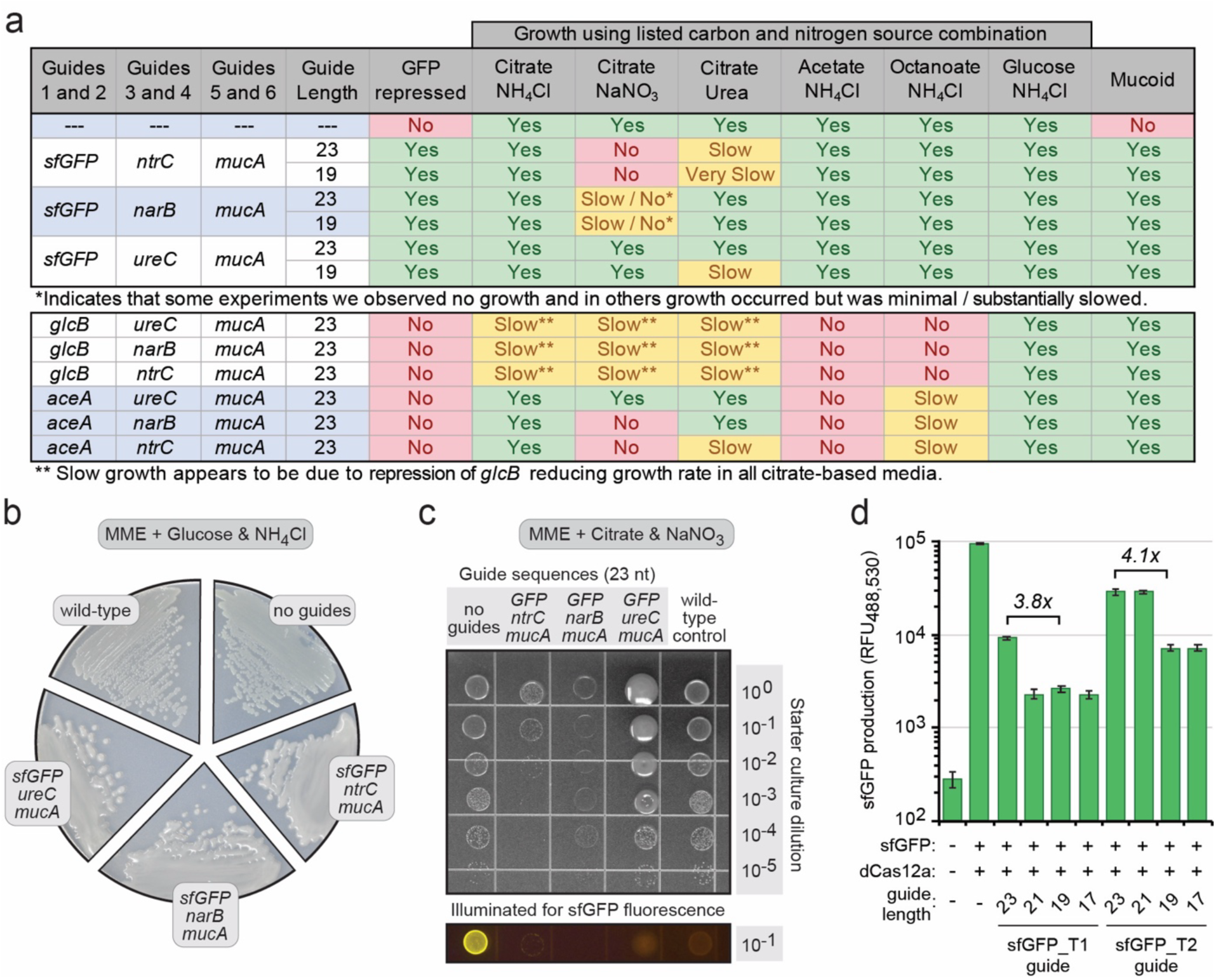
Multiplex repression of sfGFP and native pathways in *P. fluorescens*. (a) Table summarizing the phenotypes observed in multiplex gene repression assays. Results are from triplicate assays under the indicated conditions. (b) Representative data from mucoid phenotype assays. (c) Representative data from spot plate assays. Spots in each column represents 4 µL of 10-fold serial dilutions of an overnight culture grown in control medium (MME supplemented 10 mM glucose and 3 mM NH4Cl). The dilution is indicated on the Y-axis and guide sequences in the strain’s CRISPR array are indicated above. On the right is a wild-type strain lacking CRISPRi machinery. (d) Graphic displaying GFP production in *P. fluorescens* SBW25 variants as determined by flow cytometry. Presence of sfGFP, Cas12a, and composition of CRISPR array (if present) is indicated below. Error bars represent the two-sided standard deviation in 4 biological replicates.

Generally, growth was either abolished or slowed when targeting pathways involved in catabolism of carbon or nitrogen sources (**Fig. 5a**). Strains targeting *aceA* and *glcB* were unable to utilize acetate. While strains targeting *glcB* were unable to utilize octanoate, those targeting *aceA* were able to use it, albeit at a substantially lower rate than the strain with an empty array. We observed a similar pattern when targeting pathways for nitrogen sources. Strains targeting either *narB* or *ntrC* significantly inhibited nitrate utilization, but strains targeting *ntrC* or *ureC* slowed (*ntrC*) or had no apparent effect (*ureC*) on growth with urea in the tested conditions. This may be a consequence of high urease stability (*46, 47*) and expression, as observed in similar conditions in *Pseudomonas putida* (*48*). Combined, this suggests that exquisite control of gene expression may be required to repress urea catabolism.

### Truncating the 3’ end of guide sequences enhances gene repression

We next sought to evaluate whether repression could be further reinforced by truncating guide sequences from the 3’ end. A previous report demonstrated that a sfGFP-targeting guide sequence could be truncated to 16 nt before gene repression by dCas12a was impaired (*44*). So, we evaluated whether 3’ end guide truncations could improve the repression of phenotypes such as urea catabolism. First, evaluated the impact of guide truncation on sfGFP repression. For this, we truncated the 3’ end of two moderately effective sfGFP-targeting guides (sfGFP_T1 and sfGFP_T2) in 2-nt increments and evaluated the impact on sfGFP expression (**Fig. 5d**). We found that a 4-nt truncation improved repression from 10.4 to 39.8-fold (sfGFP_T1) and 3.3 to 13.6-fold (sfGFP_T2), or roughly ∼400% (**Fig. 5d**). Similar improvements were observed for both guides with 6-nt truncation, but truncation by 2-nt was inconsistent between the guides.

As guide truncation improved repression of sfGFP, we hypothesized that truncating the *ureC* and *ntrC* targeting guides will abolish use of urea as a nitrogen source. For this we generated variants of three CRISPR arrays used previously to target *sfGFP*, *narB*, *ntrC*, *ureC*, and *mucA* in which each guide was truncated by 4-nt (**Fig. 5a and Supplementary Tables S4-S5**) and evaluated the impact the truncations have on the previously measured phenotypes. Truncation of guides had no detrimental impacts on gene repressions. While truncation of *ureC* and *ntrC* guides did enable complement abolishment of growth when urea was used as a nitrogen source, the truncation of the previously ineffectual *ureC* guides substantially decreased the growth rate on urea-containing medium, and truncation of the *narB* guides further slowed growth on urea relative to strains expressing crRNAs with the full-length guides. Taken together, this data suggests that shortening the guide sequence from the 3’ end is promising method to increase CRISPRi efficacy without needing to alter Cas12a or CRISPR array expression.

### Utilizing SAGE to enables rapid implementation of CRISPRi in phylogenetically distant bacteria

SAGE enables facile reuse of genetic parts in diverse bacteria from a wide range of phyla and we demonstrate this by performing multiple reinforced CRISPRi in several additional bacteria. For this, we used SAGE to incorporate a dual fluorescent protein reporter construct into three industrially-relevant bacteria (*Pseudomonas putida*, *Rhodococcus jostii*, and *Corynebacterium glutamicum*) and an undomesticated sorghum endophyte (*Pseudomonas facilor*) (*49*) (Fig. 6a). Initially, we evaluated repression of sfGFP and mKate2 using CRISPRi expression variants that we previously optimized in *P. fluorescens* (Fig. 3). This expression variant was sufficient to repress expression of sfGFP and mKate2 to below the limit of detection in both *C. glutamicum* and the two Pseudomonads (Fig. 6a). Repression of fluorescent protein expression using this CRISPRi expression variant in *R. jostii* is modest, on par or stronger than published tools (*10, 35, 50*), and likely useful for controlling some phenotypes. The effectiveness of different guide RNAs in repressing gene expression varied widely (**Fig. 3**). Consequently, to confidently establish a correlation between phenotypes and genes of unknown function, more potent gene silencing is desirable.

**Fig. 6.**
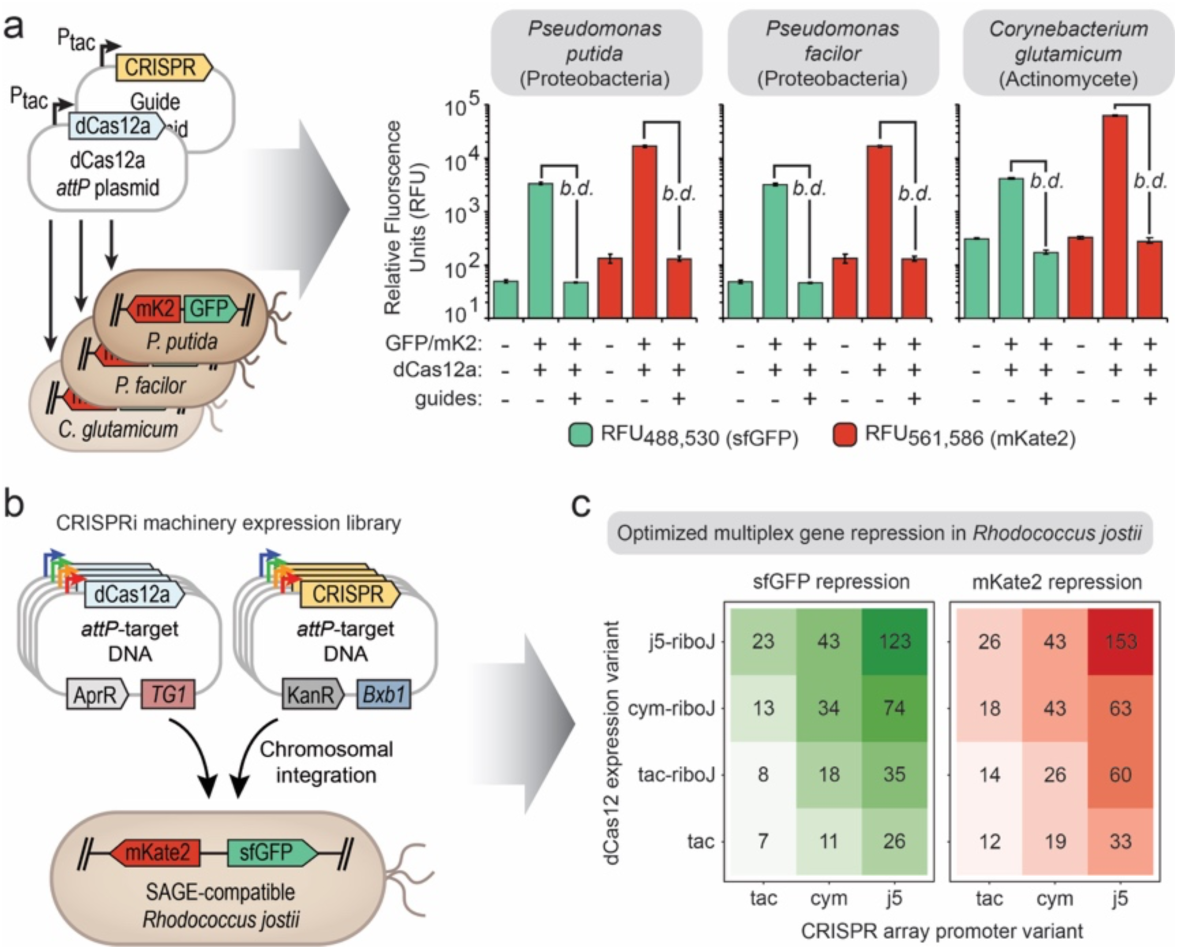
Multiplex gene repression in additional organisms. (a) Graphical representation of strain construction in multiple organisms using the same parts for CRISPRi evaluation. Graphs of sfGFP and mKate2 production as determined using flow cytometry. The organism evaluated is indicated above, presence of each fluorescent protein, dCas12a, and the composition of CRISPR arrays, fluorescence wavelength measured to detect sfGFP (green) and mKate2 (red) are indicated below. Error bars represent the two-sided standard deviation in 4 biological replicates. The notation *b.d.* indicates that fluorescence associated with protein expression was not detected (i.e., RFU is not above that of a strain that does not express the protein). (b) Graphical representation of SAGE-based combinatorial strain construction to generate CRISPRi expression variants in *R. jostii*. (c) Plot showing fold reduction in sfGFP (left) and mKate2 (right) expression relative to the fluorescent parent strain (repression). Each box represents a single strain that uses the expression variants indicated on the x- and y-axis labels. Values represent the averages of 6 biological replicates, with the exception of the strain that used tac and cym promoters for dCas12 and CRISPR array, respectively, which represents the average of 5 replicates.

Our previous work indicated that promoter strengths can vary substantially between proteobacteria and actinomycetes, and thus we sought to optimize CRISPRi in *R. jostii* by modulating the gene expression of both dCas12a and the CRISPR array. For this, we constructed a collection of expression variants that utilize both a set of strong promoters characterized in our previous work and the riboJ riboswitch (*51*), which has been shown to stabilize and increase protein expression and evaluated their performance in *R. jostii* (Fig. 6b). Increasing the expression of only a single component of the CRISPRi machinery led to at best a 3.7-fold improvement in gene repression (Fig. 6c – sfGFP repression). Highlighting the need to optimize multiple components in parallel, we found that increasing the expression of both dCas12a and the CRISPR array led to a 17.6- and 12.8-fold fold improvement in sfGFP and mKate2 repression, respectively.

## Discussion

Here we develop a CRISPRi tool that to our knowledge is the first CRISPRi toolkit that has been demonstrated to function in multiple distantly related bacterial phyla. Despite being integrated into the bacterial chromosome the gene repression observed in each organism was on par with or improved over existing CRISPRi tools others used in the same (*11, 13, 52–59*) or related genera (e.g., *Mycobacteria* (*10, 35, 50*)). For all fluorescent reporter genes evaluated in this study, gene repression increased multiplicatively when using multiple guides per gene. We were able to substantially increase CRISPRi performance in actinomycetes by evaluating an expanded the collection of dCas12a and CRISPR array expression variants. This suite of expression variants developed in this study will directly enable accelerated optimization of SAGE-based CRISPRi in diverse hosts and a similar approach could be used to generate expansive gene expression variants for other phyla.

Although there are tools that can predict the effectiveness of CRISPR guide sequences, it is still necessary to empirically test each guide to ensure that it will reliably repress genes or edit the genome. This is especially important when using a single guide to repress a gene of unknown function. Reliable gene repression is critical when evaluating the functions of conserved hypothetical or poorly annotated genes. It is important to be able to distinguish between a lack of gene disruption phenotype and a false negative arising due to insufficient gene repression. However, the inherent lack of a known phenotype for these genes means that differential mRNA or protein abundance measurements (e.g., qPCR or RNAseq) are required to empirically evaluate the effectiveness of gene repression. These measurements can be laborious and expensive, and they are not feasible when performing genome-wide functional screens.

In both this work and several other studies it has been demonstrated that using two or more guides per gene can greatly enhance gene repression (*7, 44, 50, 60*). The multiplicative increase in gene repression increases confidence that each gene will be repressed without needing to empirically validating guide efficiacy. Despite the benefits of using multiple guides per gene, most of these studies have focused on individual demonstrations of multi-guide gene repression by targeting a reporter gene in a single experiment and these results are often buried within the larger body of research (*7, 50, 60*). Furthermore, to the best of our knowledge this work is the first to demonstrate the use of multiple guides-per-gene for multiplex gene repression. We expect that by emphasizing the advantages of using multiple guides per gene the use of multiple guides per gene will become more widely adopted.

Serine recombinase-assisted genome engineering (SAGE) is a powerful technique for engineering bacteria, and here we have demonstrated that it is a valuable tool for rapidly optimizing and employing CRISPRi in diverse organisms. Chromosomal integration of genetic machinery is critical for reliable application of strains with engineered functions in complex environments and SAGE enables this in organisms that have limited high-throughput genetic toolkits. Here we use a multiplexed version of SAGE to separately incorporate the Cas protein expression cassette and CRISPR arrays into the chromosomes of bacteria from different phyla – which enables simple combinatorial assessment of Cas and CRISPR variants with minimal cloning. Previously, bacterial CRISPRi tools were developed in a boutique fashion, with each toolkit functioning in narrow ranges of organisms and multiple redundant toolkits developed for organisms in single genus (*11, 52–57*) – leading to well over a hundred published systems to-date. Here we demonstrate the application of a multiplex CRISPRi system that in principle can be used in any bacterial strain with little-to-no modifications – thus minimizing the need for researchers to expend substantial time and effort on future development of ‘boutique’ CRISPRi systems – and without the need for antibiotic selection or exogenous inducers to maintain gene repression – thus enabling it to function in environments that are suboptimal for common plasmid-based CRISPRi systems.

## Materials and Methods

### General culture conditions & media

The strains and plasmids used in this study are listed in **Supplementary Table T1**. Routine cultivation of *Escherichia coli* for plasmid construction and maintenance was performed at 37 °C using LB medium supplemented with 50 µg/mL kanamycin sulfate, 50 µg/mL apramycin sulfate, or 15 µg/mL gentamicin sulfate and 15 g/L agar (for solid medium). Cultivations for strain maintenance, competent cell preparations, and starter cultures for all Pseudomonas strains, *Rhodococcus jostii*, and *Corynebacterium glutamicum* were performed at 30 °C with 200 rpm shaking in LB (*Pseudomonas*), R2A (*Rhodococcus*), brain heart infusion – BHI (*Corynebacterium*).

MME medium (containing 9.1 mM K2HPO4, 20 mM MOPS, 10 mM NH4Cl, 4.3 mM NaCl, 0.41 mM MgSO4, 68 µM CaCl2, 1x MME trace minerals, pH adjusted to 7.0 with KOH) variants with varying carbon and nitrogen content (described below) were used in in growth assays fitness assays. 1000x MME trace mineral stock solution contains per liter, 1 mL concentrated HCl, 0.5 g Na4EDTA, 2 g FeCl3, 0.05 g each H3BO3, ZnCl2, CuCl2×2H2O, MnCl2×4H2O, (NH4)2MoO4, CoCl2×6H2O, NiCl2×6H2O.

### Plasmid construction

Q5 DNA polymerase (Thermo Scientific) and primers synthesized by Eurofins Genomics were used in all PCR amplifications for plasmid construction. OneTaq® (New England Biolabs - NEB) was used for colony PCR. Plasmids were constructed by Gibson Assembly using NEBuilder® HiFi DNA Assembly Master Mix (NEB) or ligation using T4 DNA ligase (NEB). Guide plasmids were generated by annealing multiple sets of phosphorylated oligo pairs that contained guide sequences and portions of repeat sequences and ligating them into BbsI sites of the CRISPR array containing plasmids pJE1774 or pJE1749. Plasmids were transformed into NEB 5-alpha F’I^q^ (NEB). Standard chemically competent *Escherichia coli* transformation protocols were used to construct plasmid host strains. Transformants were selected on LB (Miller) agar plates containing 50 µg/mL kanamycin sulfate, 50 µg/mL apramycin sulfate, or 15 µg/mL gentamicin sulfate for selection and incubated at 37 °C. Template DNA was isolated *P. fluorescens* SBW25 using Zymo Quick gDNA miniprep kit (Zymo Research). Zymoclean Gel DNA recovery kit (Zymo Research) was used for all DNA gel purifications. Plasmid DNA was purified from *E. coli* using GeneJet plasmid miniprep kit (ThermoScientific) or ZymoPURE plasmid midiprep kit (Zymo Research). Sequences of all plasmids were confirmed using Sanger sequencing performed by Eurofins Genomics. Ribosomal binding site variant libraries were generated using the De Novo RBS Calculator (*41*) Plasmids used in this work are listed in **Supplementary Table S1**. Maps for plasmids other than guide array variants can be found in **Supplementary File P1**. Guide sequences and arrangements can be found in **Supplementary Table T4-T5**. Oligos used for PCR screening and RBS library construction can be found in **Supplementary Table T2**.

### Strain construction

All genome modifications were performed using either the allelic exchange, integration of non-replicating plasmids into the chromosome with Serine recombinase-Assisted Genome Engineering (SAGE), or Tn insertion. Allelic exchange was performed as described previously with the exception that pK18sB was used instead of pK18mobsacB. Pairs of plasmids containing CRISPR arrays and dCas12a expression cassettes were simultaneously integrated into the chromosomes of host strains by co-electroporation with two additional non-replicating plasmids that express Bxb1 and TG1 integrases (pGW31 and pGW38, respectively). Selection for integration of both plasmids into the chromosome in *P. fluorescens* was performed using solid LB or R2A medium containing both 50 µg/mL kanamycin sulfate and 30 µg/mL gentamicin sulfate. This approach was used to both construct individual strains and expression variant libraries. Competent cells for Pseudomonads were prepared and the allelic exchange protocol used for integration of the poly-*attB* cassette into *P. fluorescens* was performed as described previously (*28, 61*). The same competent cell preparation and electroporation protocols above were used for electroporation of plasmids used for SAGE and for *in vitro* assembled EZ-Tn5 transposomes (Lucigen). The sfGFP-CmR cassette was integrated into the SBW25 chromosome by electroporating *in vitro* assembled Tn5 transposomes using EZ-Tn5 transposase (Lucigen) into SBW25 cells. DNAs for the transposomes used to incorporate the sfGFP-CmR cassette were prepared by incorporating Tn5 *att* sequences and 5’ phosphate moieties to either end of the synthetic cassette by PCR. PCR products were purified using Zymo DNA Clean & Concentrate-5 kit (Zymo Research). Transposomes containing this DNA fragment were generated *in vitro* following the manufacturer provided protocol. Two µL of each transposome assembly reaction were used for electroporation into SBW25. Selection for transformants was performed by visually identifying green fluorescent colonies. The site of transposon insertion, within the *galK* gene, was mapped using high-throughput sequencing on the MiSeq platform (Illumina) with a DNA library prepared from genomic DNA with the Nextera XT DNA library preparation kit (Illumina). A sequence map of the sfGFP cassette and insertion site can be found in **Supplementary File P1**. Allelic exchange was used to integrate a 10x poly-*attB* cassette downstream of the *ampC* gene to generate a SAGE-compatible SBW25 variant with the sfGFP-CmR expression cassette. Allelic exchange was also used to incorporate DNA cassettes containing mKate2 or mOrange2 expression cassettes into the chromosome of *P. fluorescens* SBW25 variants. Gene deletions were confirmed by colony PCR. Mutant strains and the plasmids used for each gene deletion or gene insertion are listed in **Supplementary Table T1**, respectively. Primers are listed in **Supplementary Table T2**.

Electrocompetent *R. jostii* cells were prepared by pre-culturing at 30 °C in LB until cultures reached stationary phase. Fresh medium was inoculated with the preculture to an OD600 of 0.01 and incubated aerobically until the cells reached mid-log phase (∼OD600 = 0.4 - 0.6). Once desired culture density was reached, the cells were centrifuged at 5000 x g for 20 min at 4 °C and washed at ½ culture volume ice-cold 10% glycerol. Centrifugation and washing was repeated 2 additional times, and then cells were resuspended in 1/50^th^ culture volume ice-cold 10% glycerol. Competent cells were either used immediately or stored at -80 °C. Electroporation was performed as follows. DNA was added to 70 µL ice-cold electrocompetent cells and incubated on ice for 30 minutes. Following incubation competent cells were transferred into an ice-cold 0.1 cm gap electroporation cuvette and electroporated at 1.6 kV, 25 µF, and 200 ohms. Following electroporation, 950 µL of R2A was added to the competent cells and the resulting mixture was incubated aerobically, at 30 °C with shaking for 3 hours. Following this recovery step, various dilutions of recovery cultures were plated on selective R2A solid medium and incubated aerobically at 30 °C. Colonies typically appear after ∼2 days when *R. jostii* is grown on solid medium.

Electrocompetent *C. glutamicum* cells were prepared by pre-culturing at 30 °C in 25 mL of BHIS for 24 hours in a 125 mL shake flask. The cells were centrifuged at 4000 rpm for 10 minutes and resuspended in 1x MME salts. In a 250 mL flask, 50 mL of NCM medium was inoculated with the resuspended preculture to an OD600 of 0.3 and incubated aerobically at 30 °C for approximately 3-6 hours. Once the NCM incubation step was complete, the culture was chilled on ice for 10 minutes and centrifuged at 4000 rpm for 10 minutes at 4 °C and washed with ½ culture volume ice-cold 10% glycerol. The cultures remained on ice and centrifugation and washing were repeated 2 additional times. Following the washes, the cells were resuspended in 1/50*th* culture volume ice-cold 10% glycerol. Competent cells were either stored at -80 °C or used immediately in a subsequent electroporation. An 80 µl aliquot of ice-cold competent cells was incubated on ice with DNA for 30 minutes. After incubating, competent cells were added to an ice-cold 0.2 cm gap electroporation cuvette and electroporated at 2.5 kV, 600 ohms, and 25 uF. BHI medium was pre-warmed at 55°C and 920 µL was added to the electroporated competent cells and incubated aerobically, at 30 °C with shaking for 2-4 hours. Following recovery, the cells were plated on selective BHI solid agar plates and incubated aerobically at 30 °C. Colonies typically appear after 1-2 days when *C. glutamicum* is grown on solid medium. BHIS is a modified BHI medium containing 91.1 g/L sorbitol. NCM medium contains 91.1 g/L sorbitol, 17.4 g/L K2HPO4, 11.6 g/L NaCl, 5 g/L glucose, 5 g/L Tryptone, 3 g/L glycine, 1 g/L DL-threonine, 1 g/L yeast extract, 0.2 g/L trisodium citrate, 0.05 g/L MgSO4- 7H2O, 0.4% isoniazid (INH), 0.1% Tween 80 and its pH is adjusted to 7.2 prior to filter-sterilization.

For evaluating CRISPRi in organisms other than *P. fluorescens* we performed the following. The dual fluorescent marker cassette (sfGFP/mKate2) containing plasmid pAF26 was integrated by integrated into the R4 *attB* by co-electroporation with a non-replicating plasmid that expresses the R4 integrase (pGW39). The CRISPR array and dCas12a plasmids were integrated by co-electroporation with plasmids expressing the Bxb1 integrase (pGW31) and TG1 integrase (pGW38), respectively. Selection for integration was performed using solid LB (*P. putida* and *P. facilor*), R2A (*R. jostii*), or BHI (*C. glutamicum*) containing the following antibiotics: Fluorescent marker plasmid – 30 µg/mL (Pseudomonads and *R. jostii*) or 10 µg/mL (*C. glutamicum*) gentamicin sulfate; CRISPR array plasmids – 50 µg/mL kanamycin sulfate (all organisms); dCas12 plasmids – 50 µg/mL (Pseudomonads and *C. glutamicum*) or 30 µg/mL (*R. jostii*) apramycin sulfate. After the initial chromosomal insertion selection is no longer necessary.

### Flow cytometry assessment of fluorescent protein repression

Starter cultures for fluorescent protein repression assays were prepared by cultivating cells from either glycerol stocks or transformant colonies in 1mL LB, R2A, or BHI in 96-well deep-well plates at 30 °C with 1200 rpm shaking (1 mM orbital) until they reached stationary phase. At least three replicate assay cultures for each strain were prepared by diluting the overnight cultures 1:100 into fresh media in another 96-well deep well plate containing fresh media and incubating until cultures reached stationary phase. Stationary phase cultures were assessed for construct stability by flow cytometry. Briefly, stationary phase cultures were diluted 200-fold in 1x phosphate buffered saline (PBS) and fluorescence was measured using a NovoCyte Flow Cytometer (Acea Biosciences). Green fluorescence (sfGFP) was measured using a 488 nm laser with a 530/30 nm filtered detector. Red fluorescence (mKate2) was measured using a 561 nm laser with a 586/20 nm filtered detector. A preliminary gating of 2500 and 10 were used for FSC and SSC, respectively, and at least 25,000 events were measured for each sample. Median relative fluorescence (RFU) values were used for fluorescent protein expression calculations. For calculation of fold-repression the background autofluorescence of a strain lacking the fluorescent protein cassette was subtracted from the median of all samples. Flow cytometry-based measurements was used for all experiments other than those displayed in **Fig. 4c-d** and **Supplementary Fig. S2** – which use mOrange2 in *P. fluorescens* SBW25.

### Microtiter plate reader assessment of fluorescent protein repression

Microtiter plate reader-based assays were used to measure fluorescent protein production in strains that contained mOrange2 – which are displayed in Fig. 4c-d and Supplementary Fig. S2. While the laser and filter combinations used with the Novocyte flow cytometer distinguish green versus red fluorescent proteins, the broad bandwidth of the filters do not allow the instrument to distinguish individual contributions of mNeonGreen, mOrange2 and mKate2 to cell fluorescence. So, we used a BioTek Neo2SM plate reader that is equipped with a monochromator with 1 nm bandwidth for both emission and excitation – which enables the contributions of the three proteins toward fluorescence to be resolved.

Cultures were grown as described for flow cytometry-based experiments. Once in stationary phase, aliquots of each culture were diluted in LB such that the OD600 measurements are the linear range of measurement for the plate reader. This was typically 4-fold. The green, orange, and read fluorescence values were measured with ex501/em520, ex545/em570, and ex588/em630 respectively. Bandwidths of 9, 12, and 20 nm were used for green, orange, and red measurements, respectively. Fluorescent protein expression was calculated using RFU/OD600 and autofluorescence background from a strain lacking fluorescent proteins was subtracted from these values to determine fold-repression.

### Visible phenotype evaluation assays

*P. fluorescens* strains with triple gene knockdown CRISPR arrays, non-targeting CRISPR arrays, or no CRISPR-Cas system were revived from glycerol stocks in by overnight cultivation in LB at 30 °C. Cultures were diluted 100-fold into MME medium containing 10 mM glucose and 3 mM NH4Cl as carbon and nitrogen sources, respectively, and incubated until they reached stationary phase (typically 24 hrs). Subsequently, cultures were serially diluted 7 times by 10-fold, and 5 µL of each dilution was spotted onto solid MME containing either 10 mM glucose, 10 mM sodium citrate, 30 mM sodium acetate, or 7.5 mM sodium octanoate as carbon sources and either 5 mM ammonium sulfate, 10 mM sodium nitrate, or 5 mM urea as nitrogen sources. Additionally, cultures were spread onto solid MME with 10 mM glucose and 5 mM ammonium sulfate and dilutions spread onto solid LB with and without 5% sucrose to evaluate mucoid phenotypes. All plates were incubated at 30 °C and imaged with a Nikon D7500 dSLR camera equipped with a AF-S DX NIKKOR 18-300mm f/3.5-6.3G ED VR Lens when the largest colonies were readily visible by the eye. Images were taken with white light or using a Dark Reader blue light box (Clare Chemical) with orange filter to capture green fluorescence.

## Acknowledgements

This research was supported in part by the U.S. Department of Energy (DOE), Office of Biological and Environmental Research (BER), as part of BER’s Genomic Science Program (GSP), and is a contribution of the Pacific Northwest National Laboratory (PNNL) Secure Biosystems Design Science Focus Area “Persistence Control of Engineered Functions in Complex Soil Microbiomes”. Additional support was provided by the Laboratory Directed Research & Development program at PNNL. PNNL is a multi-program national laboratory operated by Battelle for the DOE under Contract DE-AC05-76RL01830. We also thank Adam Guss for kindly sharing SAGE-compatible *Pseudomonas putida* (AG5577) and *Corynebacterium glutamicum* (AG6216) strains.

