## Supplementary Figs S1-S3 for "Reinforced CRISPR interference enables reliable multiplex gene repression in phylogenetically distant bacteria"

**Supplementary Figures**

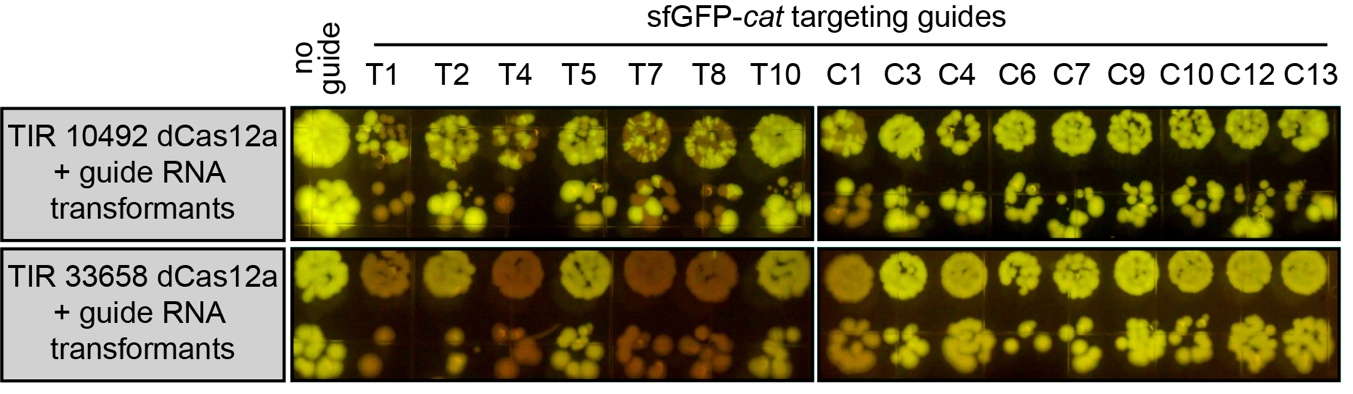

**Supplementary Fig. S1.** Images of SBW25 transformants that express the indicated Cas12a variant (left) and contain a CRISPR array containing the guide sequence indicated above. Images data for ‘no guide’ and template strand targeting guides (T#) are also found in Fig 2 and are included here for visual comparison with coding strand-targeting guides (C#).

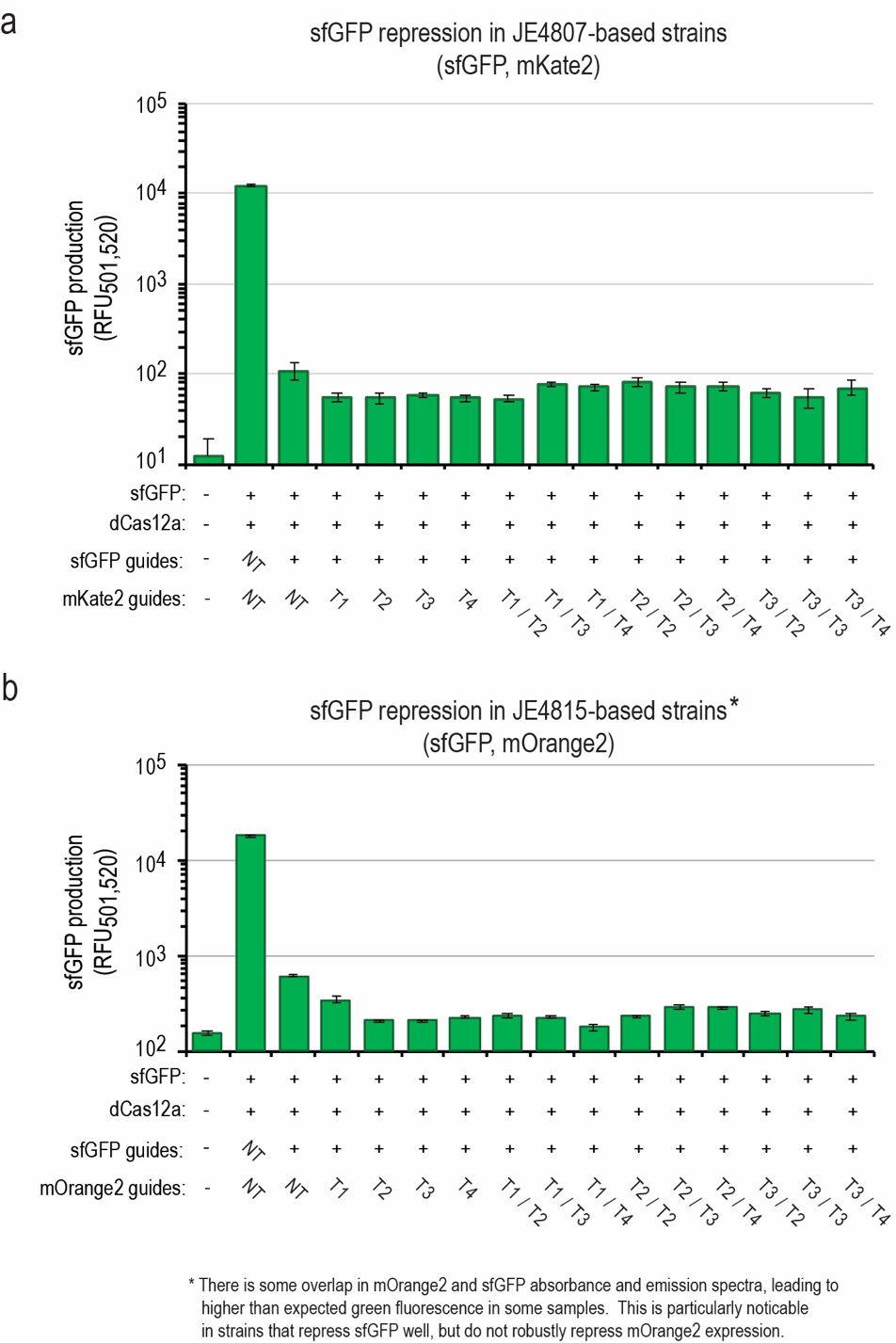

**Supplementary Fig. S2.** Graph displaying production of sfGFP in P. fluorescens SBW25 strains that express either (a) sfGFP and mKate2 or (b) sfGFP and mOrange2. Presence of fluorescent protein, Cas12a, and composition of CRISPR arrays are listed below. Y-axis values represent either (a) the mean fluorescence as determined using flow cytometry or (b) plate reader measurements. Error bars represent the two-sided standard deviation in 4 biological replicates. *There is some overlap in mOrange2 and sfGFP absorbance and emission spectra, leading to green fluorescence that is attributable to mOrange2. This is particularly noticeable in strains that exhibit robust sfGFP repression and poor-to-no repression of mOrange2.

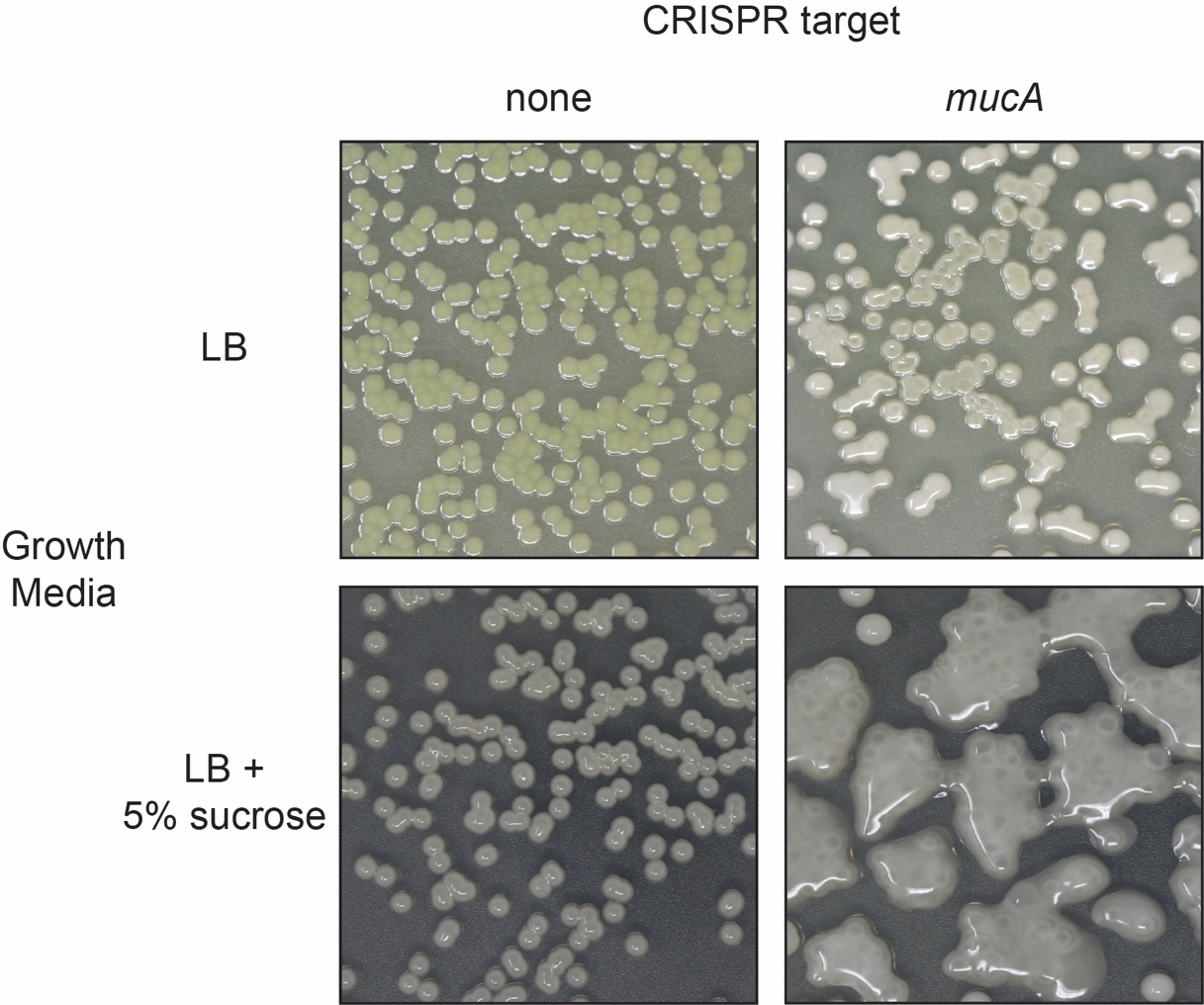

**Supplementary Fig. S3.** Photographs of P. fluorescens JE4694 strains cultivated on solid agar media to evaluate mucoid phenotypes. Images were taken after ~24 hours of incubation. Both strains contain the dCas12a expression cassette. The strain on the left panels (none) contains a CRISPR array that has no guides targeting the mucA gene, while the strain on the right panels contains a CRISPR array with two guides targeting the coding region of mucA.
